# Dissociable Effects of Curiosity and Hedonic Valence on Reinforcement Learning

**DOI:** 10.1101/2025.06.24.660807

**Authors:** Kathryn M. Rothenhoefer, McKenna D. Romac, Krystal Henderson, Vincent D. Costa

## Abstract

Curiosity and exploration support learning and adaptive decision-making in uncertain environments. While these processes are sensitive to motivational context, it remains unclear how outcome valence shapes exploration across species. Human studies suggest that aversive contexts increase exploration, but these effects often rely on verbal framing and explicit instructions. To gain deeper insight into how exploration strategies emerge from experience alone, this study investigated the influence of hedonic valence on novelty seeking, exploration, and reinforcement learning in rhesus macaques. Using visual tokens as secondary reinforcers, we found that monkeys explored novel, uncertain options more frequently when exploitation would lead to losses rather than gains. However, our analyses clarified that this heightened novelty seeking was primarily a consequence of the monkeys employing an optimistic prior belief about the value of novelty, rather than a categorical, valence-dependent shift in their underlying curiosity or the information bonus associated with exploration. Approach and avoidance motivation did influence other aspects of reinforcement learning. Monkeys demonstrated faster learning from losses than from gains, indicating that they were averse to losing tokens. They also frequently chose an option and then quickly aborted their choice. These choice balks were strategic responses to approach–avoidance conflicts and uncertainty, and represented self-generated bouts of exploratory behavior that led to valence-dependent use of directed and random exploration. These findings suggest that different strategies are used to manage explore–exploit tradeoffs induced by novelty or internal motivational conflicts, revealing dissociable effects of curiosity and hedonic valence on reinforcement learning.

---

The explore-exploit dilemma refers to the difficulty of balancing exploration of novel choice opportunities with exploitation of known rewards (Cohen et al., 2007; Wilson et al., 2021). Novelty seeking—a widely observed solution to this dilemma across species—enables humans and animals to learn whether future actions could yield greater rewards (Costa et al., 2014; Wittmann et al., 2008), and is a core component of curiosity (Bradley, 2009; Kidd & Hayden, 2015; Monosov, 2024). However, the majority of what is known about the psychology and neuroscience of explore-exploit decision making focuses on scenarios in which exploration resolves uncertainty about exclusively appetitive outcomes (Wilson et al., 2021). In naturalistic environments, exploration also carries the risk of discovering aversive outcomes. This tension between motivational goals to approach or avoid, and tradeoffs between exploring or exploiting to maximize the utility of future actions, highlights the need to understand how hedonic valence shapes exploratory decision making.

Prior studies of explore-exploit decision making in humans have exclusively examined decisions to explore or exploit under separate motivational goals: maximize appetitive outcomes or minimize aversive outcomes. In multi-armed bandit tasks humans overexplore uncertain options when choices result in different size losses compared to how often they explore when choices result in different size gains (Krueger et al., 2017; Lejarraga & Hertwig, 2017; Wilson et al., 2014). These biases are observed in healthy participants, suggesting that overexploration in aversive environments may serve as an adaptive heuristic. Consistent with this view, explicit anxiety inductions reduce exploration in healthy participants (Li et al., 2025). However, prior studies of explore-exploit decision-making in clinical populations have not revealed a transdiagnostic pattern of over- or underexploitation that would support this perspective (Brown et al., 2023). Rather, patients diagnosed with mood disorders exhibit difficulties in discerning irreducible from unexpected uncertainty, leading to disruptions in reinforcement learning that produce a variety of imbalances in explore-exploit decision making (Brown et al., 2023). Understanding how these imbalances relate to the different strategies that can be used to resolve the explore-exploit dilemma is at the crux of developing explore-exploit decision making as a paradigm relevant to the goals of computational psychiatry (Lloyd et al., 2024).

Decisions to explore are prompted either by external changes in the decision environment—such as the introduction of novel or uncertain choice options in multi-armed bandit (Costa & Averbeck, 2020; Costa et al., 2014, 2019; Hogeveen, Mullins, et al., 2022) or horizon tasks (Krueger et al., 2017; Wilson et al., 2014) — or by internal changes in beliefs about the value of existing choice options (Ebitz et al., 2018). Two main strategies are observed: directed or random exploration. Directed exploration is when decision makers forego known rewards in order to sample an uncertain option to gain information (Wilson et al., 2014). Random exploration describes off-policy choices that do not necessarily provide additional information (Wilson et al., 2014). It has been shown that in both appetitive and aversive decision environments that human decision makers use directed and random exploration to solve the explore-exploit dilemma, and that directed exploration is heightened in aversive choice environments (Krueger et al., 2017).

Optimal strategies for solving the explore-exploit dilemma apply when outcomes are purely appetitive or aversive (Averbeck, 2015), but directed exploration becomes more difficult when outcome valence is uncertain. One solution is to weigh information gained through exploration differently based on whether exploitation leads to appetitive or aversive outcomes. According to the Bayesian shrinkage hypothesis (Krueger et al., 2017), value estimates are biased toward optimistic or pessimistic priors depending on the valence, uncertainty, and magnitude of choice outcomes. When uncertainty is high and decision makers lack information regarding the valence of choice outcomes, they rely more heavily on these priors to guide explore-exploit decisions. However, translating these effects into nonverbal animal models has been challenging. For example, well-established reward horizon effects (Krueger et al., 2017; Wilson et al., 2014, 2021) on explore-exploit decision making in humans are not replicated in rodents (Wang et al., 2023) or monkeys (Jahn et al., 2023).

Therefore, developing reinforcement learning tasks that can be used across species and that equate appetitive and aversive outcomes to test whether approach and avoidance motivations differently shape exploratory decisions remains a key challenge. This is especially important for two reasons: First, motivation and affect are now recognized as core components — not just modulators — of decision making (Lerner et al., 2015; Lopez-Guzman & Atlas, 2024; Shenhav, 2024). Second, forward and backward translation of findings across model systems is needed as it becomes clear that the neural systems mediating explore-exploit decision making are not solely governed by prefrontal cortical circuits unique to primates (Cohen et al., 2007; Daw et al., 2006; Hogeveen, Medalla, et al., 2022; Mansouri et al., 2017), but also evolutionary conserved subcortical motivational circuits (Ahmadlou et al., 2025; Averbeck & Costa, 2017; Costa et al., 2019; Monosov, 2024).

Token economies provide a powerful tool for examining shared psychological and neural mechanisms of decision-making across species (Ayllon & Azrin, 1965; Cowles, 1937; DeFulio et al., 2014; Hackenberg, 2018; Kelleher, 1958; Phillips, 1968; Wolfe, 1936). Previous work has shown that monkeys can learn to discriminate between cues predicting a gains or losses of virtual tokens, which are later exchanged for primary rewards (Banaie Boroujeni et al., 2022; Burk et al., 2024; Seo & Lee, 2009; Taswell et al., 2023; Yang et al., 2022). Unlike primary reinforcers—which differ in sensory characteristics and salience (Lang & Bradley, 2010; Miller, 1959)—secondary reinforcers can be explicitly quantified in a common motivational currency. This allows more precise comparisons of approach and avoidance motivation during exploratory decision making and can reveal novel strategies not typically observed when only primary reinforcers are used.

While human studies have examined exploration in appetitive and aversive contexts, prior studies of explore-exploit decision making in nonhuman primates have exclusively examined exploration in the context of appetitive, primary reinforcers (Costa & Averbeck, 2020; Costa et al., 2014, 2019; Ebitz et al., 2018; Giarrocco et al., 2024; Hogeveen, Mullins, et al., 2022). A major barrier to studying exploration in aversive contexts in monkeys is the difficulty of finding appetitive and aversive outcomes that elicit equally strong approach and avoidance behaviors (Amemori & Graybiel, 2012; Amemori et al., 2015; Kobayashi et al., 2006; Leathers & Olson, 2017; Roesch & Olson, 2004). To address this, we developed a token-based economy where tokens—exchangeable for juice—served as a common currency in both gain and loss contexts. This enabled us to study novelty-driven exploration when outcomes could be either appetitive or aversive.

We found that monkeys’ decisions to explore or exploit were not influenced by the valence of known outcomes, but by their optimistic valuation of novelty. Like humans (Krueger et al., 2017), monkeys used Bayesian shrinkage to guide directed exploration. However, approach and avoidance motivation did influence other aspects of reinforcement learning. Monkeys learned faster from losses than from gains and differentially deployed directed and random exploration to resolve approach–avoidance conflicts. These findings clarify how decision strategies are shaped by outcome valence and show how primates manage uncertainty when exploration can lead to either appetitive or aversive outcomes.

## Results

### Secondary Reinforcers Enable Assessment of Valence-Dependent Reinforcement Learning

Two rhesus monkeys performed a three-arm bandit task with the goal to acquire visual tokens that were exchanged for primary rewards (**Figure 1A**). The monkeys were trained prior to the task to associate the exchange of individual tokens with receipt of a fixed amount of a primary juice reward (see Methods). The monkeys began each session with a fixed token endowment of eight tokens. Each trial started by displaying the visual tokens the monkey had accumulated at the bottom of the screen. Next, the monkeys had to acquire and hold central fixation for a variable duration (500-750 ms). If central fixation was completed, this triggered the presentation of three visual cues. If central fixation was not completed, the trial was aborted and restarted without the monkey knowing what choices were available (central fixation abort; **Figure 1A**). Following presentation of the three visual cues, the monkey indicated its choice by completing a saccade and fixating one of the cues (500 ms). If the monkey did not make a saccade, or did not successfully hold fixation on a cue (choice balks), the trial was aborted and restarted with the monkey now knowing what choices were available (**Figure 1A**). Each cue was associated with a reward distribution that determined the number of tokens gained or lost, following its selection. Each choice outcome was signaled by updating the visual token count at the bottom of the screen. If a choice resulted in the loss of all the accrued tokens, no tokens were displayed. Token counts could not fall below zero and no visual tokens were displayed when outcomes would otherwise lead to a token debt (e.g., selecting a loss cue when starting the trial with 0 tokens). The number of visual tokens displayed at the end of the trial determined the probability of whether the tokens were exchanged for primary rewards (i.e., cashed out; **Figure 1D**). If the visual tokens were cashed out, the displayed tokens disappeared from the screen one by one immediately followed by receipt by a fixed quantity of apple juice. The token endowment was then refreshed with eight tokens and the next trial started after a fixed length intertrial interval (ITI). If the tokens were not cashed out, the monkey experienced the same length ITI. The accumulated tokens carried over and were then displayed at the start of the next trial.

**Figure 1.**
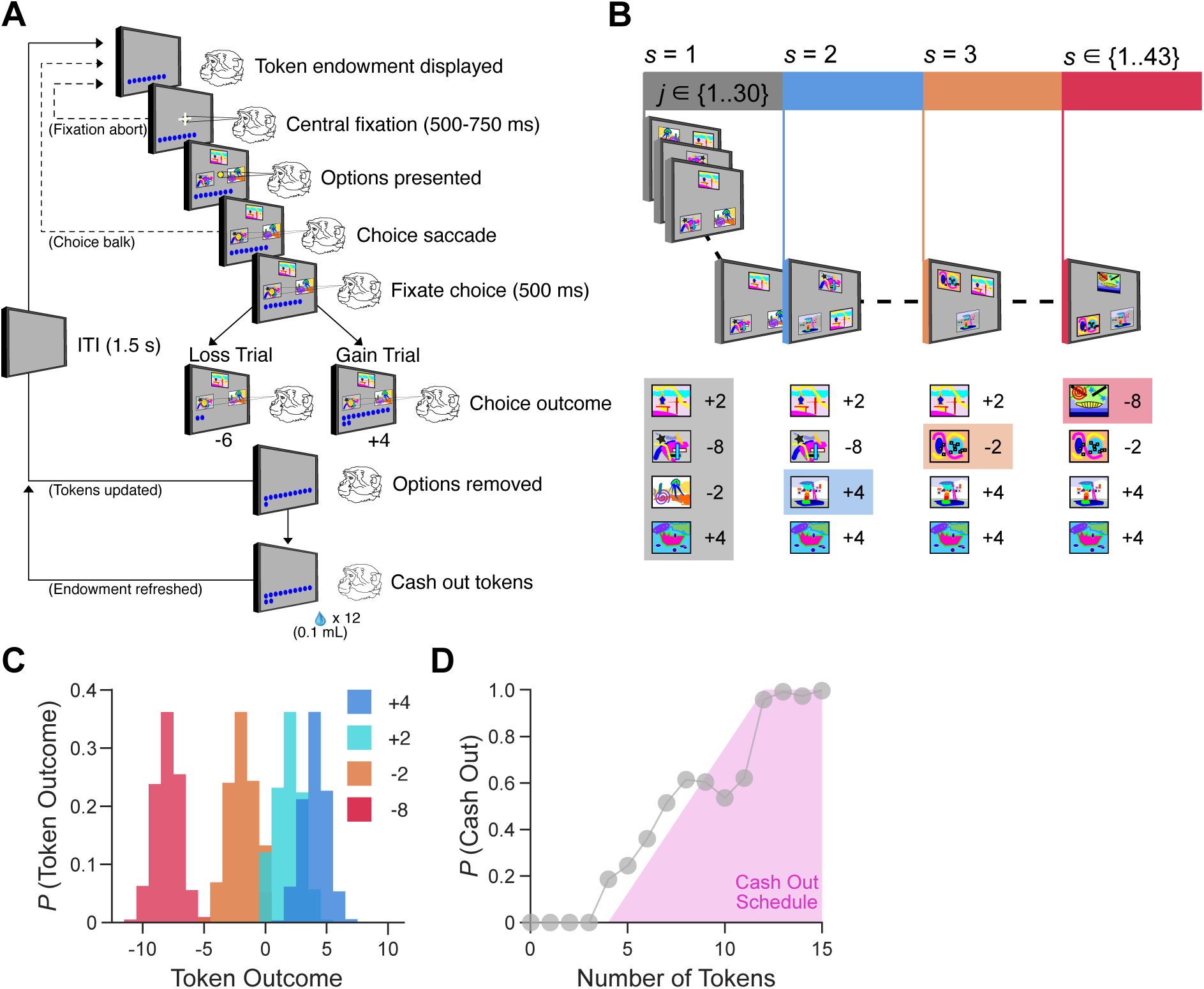
Three-arm bandit task that induces explore-exploit tradeoffs via novel options associated with appetitive and aversive outcomes. **(A)** Structure of an individual trial in the three-arm bandit task. **(B)** Each block (s) of trials (j), up to a total of 800 trials. Each block contained a minimum of 10 and a maximum of 30 trials. On each trial, three visual images were randomly selected from a set of four images specific to each block. The start of a new block was indicated by one of the four cues being replaced by a novel image, such that in the new block there are now three known alternative cues carried over from the previous block and one novel cue. The new cue can take on any value, regardless of the values of the alternative options. **(C)** Each cue was assigned an outcome distribution centered on a mean of either +4, +2, −2, or −8 tokens (SD = 1). The actual outcome following the selection of a cue was randomly selected from a normal distribution surrounding the average token amount. The token outcome selected was always the same valence as the average token amount, for example, the minimum outcome that could happen following selecting a +2 cue was 0 tokens. **(D)** Experienced probability of cashing out their tokens by the number of tokens at the end of a trial (gray), and the programmed cash out schedule (pink).

We induced explore-exploit tradeoffs by replacing an option whose value the monkey already learned with a novel option whose value was uncertain (**Figure 1B**). This occurred every 10 to 30 trials. During each block of trials, the monkeys learned to choose between four cues, each assigned one of four independent reward distribution schedules (**Figure 1C**). On each trial, three of the four cues were randomly selected and presented to the monkey. This was done to limit the monkeys’ anticipation of choosing a particular option from trial-to-trial. To learn the value of the novel option, the monkeys had to explore, foregoing the opportunity to exploit previously learned options. Based on the outcome distributions assigned to each cue, choice outcomes were drawn from one of four distributions that had an average value of either −8, −2, +2, or +4 tokens (**Figure 1C**). Reward distributions were clipped so that gain and loss cues were exclusively associated with appetitive and aversive outcomes. For example, a cue with an *a priori* value of +2 had a minimum token outcome of 0 tokens, and a cue with an *a priori* value of −2 had a maximum outcome of 0 tokens. The large loss token outcome distribution was −8 instead of matching the large gain cue of +4 to increase the monkeys’ motivation to discriminate between small and large loss cues, as previous work has shown monkeys will not always readily discriminate between different value losses (Taswell et al., 2018). Finally, the cash out schedule was programmed so that the probability of the accrued tokens being cashed out was zero for up to four tokens accrued and then increased linearly (.125 step in probability) between five and 12 tokens (pink shaded region in **Figure 1D**). To address within-session shifts in motivation, the window in which the monkeys could not get a token cashout was shifted up or down, resulting in slight deviations from the programmed cash out probabilities (gray line in **Figure 1D**). The monkeys could not have their tokens cashed out if they had received a cash out on the previous trial.

### Value Rather Than Valence Influences Novelty Seeking

Since all options were novel at one point and were independently assigned a distribution of token gains or losses, we assessed if the monkeys were efficient in learning about the valence and value of novel options (**Figure 2A**). Averaging the monkeys’ choices over the first 20 trials since a novel option was introduced showed that the monkeys learned to discriminate the novel options based on their *a priori* value assignments (Value: *F*(3, 315) = 289.99, *p* < .001). The monkeys were more likely to choose cues associated with large compared to small gains (*F*(1, 105) = 33.25, *p* < .001) and were more likely to choose cues associated with small compared to large losses (*F*(1, 105) = 45.22, *p* < .001).

**Figure 2.**
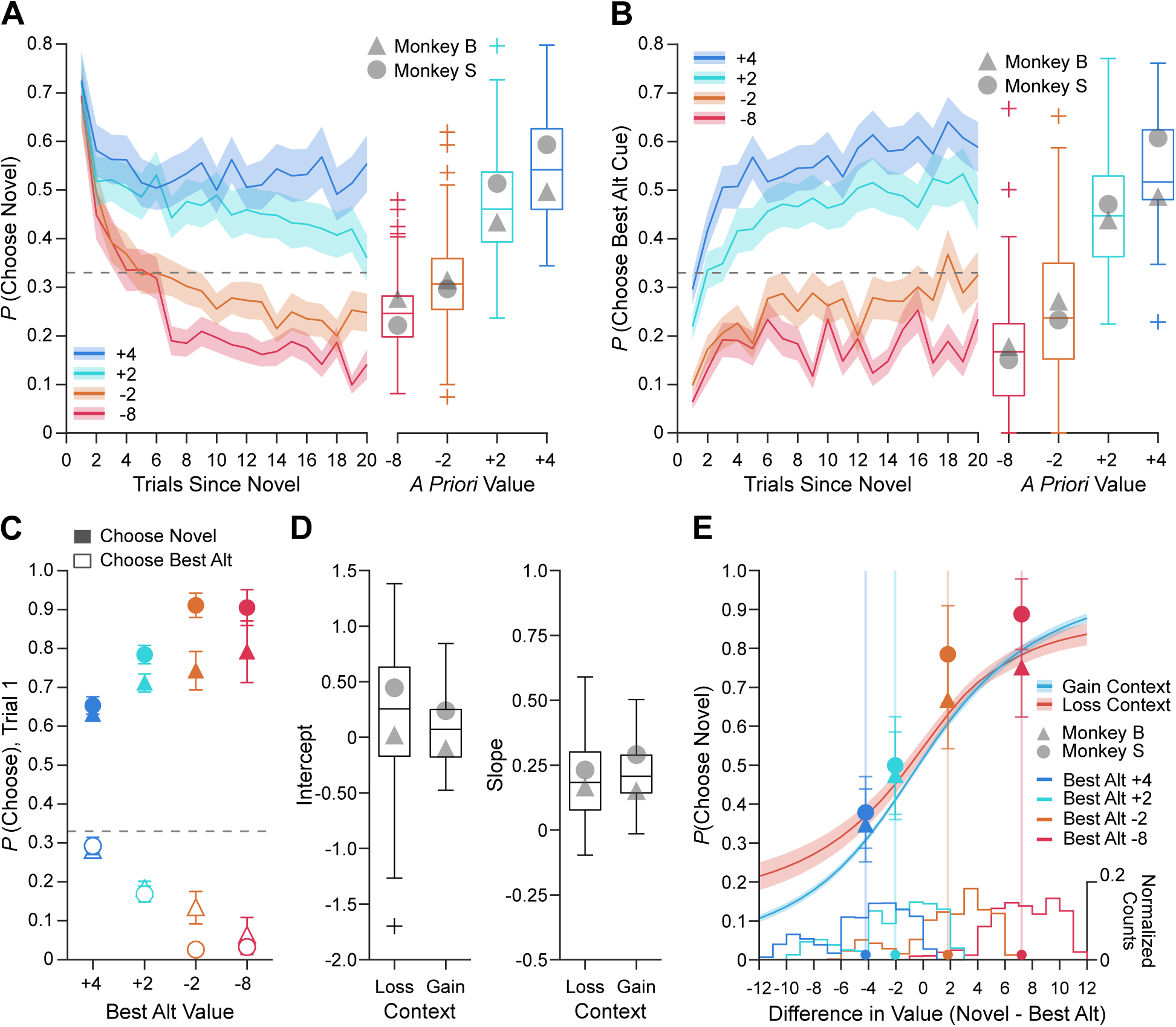
Value differences and not hedonic valence influences novelty seeking. **(A)** Probability of choosing the novel option as a function of trials since the novel cue was introduced, separated by the *a priori* value of the novel cue. Box plots include data across 20 trials. Gray markers are the individual monkeys’ averages. Grey dashed line is 33%, or chance choice probability across the three available cues, for panels **A**, **B** and **C**. **(B)** Probability of choosing the best alternative option as a function of trials since the novel cue was introduced, separated by the *a priori* value of the best alternative cue. Box plots include data across 20 trials. Gray markers are the individual monkeys’ averages. **(C)** Probability of choosing the novel option and the best alternative option on the first trial that the novel option is introduced, separated by the *a priori* value of the best alternative cue. Triangle markers represent Monkey B’s averages while circle markers represent Monkey S’s averages. Filled markers represent the probability of choosing the novel option and open markers represent the probability of choosing the best alternative option. Error bars are S.E.M across sessions. **(D)** Fitted intercept (left panel) and slope (right panel) coefficients from logistic regressions run on session-by-session data in choice situations where the best alternative option was a loss cue or a gain cue (Loss vs. Gain Context). Grey markers are the individual monkeys’ average coefficients across sessions. **(E)** Probability of choosing the novel option as a function of the difference in empirical value between the novel option and the best alternative option, estimated from the average intercept and slope for gain (Blue) and loss (Red) contexts. Shaded regions are S.E.M. across session estimates using session-by-session slope and intercepts for each context, averaged across monkeys. The second axis represents the normalized counts of value differences experienced for each possible *a priori* value of the best alternative cue. These bin counts were estimated within a session, averaged across sessions within monkeys, and then averaged across monkeys. Filled circles along the x-axis and the lines emanating from them show the average empirical value difference (Novel - Best Alt) experienced for each *a priori* value of the best alternative cue. The markers on those lines represent each monkeys’ median probability of choosing the novel option when the value difference between the novel and the best alternative option was +/− 0.5 of the average empirical value difference, for each of the four *a priori* values that the best alternative option could take. Error bars are S.E.M across sessions.

Exploration of novel options allowed the monkeys to resolve uncertainty and learn both the valence and value of each option. To further characterize the monkeys’ ability to exploit what they learned, we examined how frequently they chose the best alternative option, based on its assigned value (**Figure 2B**). There was a clear preference for the best alternative option when it led to a gain in tokens instead of a loss in tokens (Value: *F*(3, 280) = 239.74, *p* < .001). Similar to the novel options, cues associated with losing tokens were routinely avoided, even when they represented the best alternative option, and monkeys showed a preference for cues that led to small compared to large losses (*F*(1,105) = 26.96, *p* < .001). Additionally, the monkeys displayed a stable preference for options associated with large compared to small gains (*F*(1, 70) = 69.49, *p* < .001).

The monkeys’ increased selection of novel, uncertain options on the first trial when the best alternative is associated with losing compared to gaining tokens implies valence-dependent changes in exploration. To test this hypothesis, we reexamined how frequently monkeys selected the novel and the best alternative option on the first trial it was introduced, based on the *a priori* value of the best alternative option (**Figure 2C**). Despite being unable to predict the valence or specific value assigned to a novel option when it was first introduced and prior to receiving any feedback, the difference in monkeys’ selection of the novel vs. the best alternative option increased as the value of the best alternative option decreased (Value x Choice: *F*(3, 251) = 28.5, *p* < .001). This result aligns with prior studies in humans, demonstrating increased directed exploration in loss contexts. This contrasted with the monkeys’ selection of the worst alternative option, which did not vary based on the value of the best alternative option (Value: *F*(3, 251) = 0.42, *p* = .736). Selection of the worst alternative option is a proxy for random exploration.

Unlike prior studies, however, novelty was equally associated with gains and losses in our task. It was therefore not surprising that the monkeys preferred the novel option when the best alternative option was associated with losing tokens. In these choice scenarios, the mean empirical value of the novel option (*M* = 0.19, *SD* = 2.68) was higher than the average best alternative option in loss contexts (*M* = −2.63, *SD* = 1.41). To determine whether monkeys were more exploratory in aversive contexts or whether they were simply choosing to explore in line with differences in expected value, we computed the difference in the empirical value between the novel and best alternative options for all trials, and separately fit logistic regression models to predict the monkeys’ selection of the novel option when the best alternative options were associated with losing (aversive context) or gaining tokens (appetitive context). The intercept of the logistic regression function fit across sessions quantified the monkeys’ overall novelty preference and magnitude of the information bonus associated with exploring. On average, that information bonus did not differ based on the valence context (left panel of **Figure 2D**; *F*(1, 105) = 0.02, *p* = .90). The slope of the fitted regression functions quantified how the monkeys’ novelty preference changed as a function of the trial-by-trial empirical value difference between novel and the best alternative option. On average, the fitted slope did not differ by the valence of the best alternative option (right panel of **Figure 2D**; *F*(1, 105) = 0.08, *p* = .78). The lack of an effect of valence and a clear effect of value on exploratory decision making is apparent when the predicted likelihood that the monkeys chose to explore novelty based on the difference in value between the novel and best alternative options is plotted against the monkeys’ actual choice behavior (**Figure 2E**). Therefore, heightened novelty seeking in loss contexts is due to the value differences rather than a categorical, valence-dependent shift in novelty valuation.

### Enhanced Learning from Losses Compared to Gains

We next asked whether learning dynamics differed by outcome valence. The monkeys’ choices indicated they learned, within each valence category, to discriminate between cues associated with small and large gains or losses (**Figure 2A** and **B**). To test whether monkeys were more sensitive to losses compared to gains in our task, and whether they learned aversive associations faster than appetitive associations, we fit a reinforcement learning model to the monkeys’ choice behavior. The best-fitting reinforcement learning model included three parameters that accounted for the monkeys’ explore-exploit tendencies: a choice consistency term (**Figure 3F**), a novelty bonus (**Figure 3G**), and a perceptual novelty discount (**Figure 3H**) (Costa et al., 2014, 2019; Kakade & Dayan, 2002; Wittmann et al., 2008). This model performed better than a related model that did not include a perceptual novelty discounting term (*χ^2^* = 11.66, *p* = .0006). The estimated novelty bonus indicated the monkeys assigned a high initial value to novel options before they were chosen (**Figure 3G**). An alternate model specifying separate bonuses for when novel options were introduced in appetitive or aversive contexts did not improve model fit (*χ^2^* = 1.58, *p* = .2083). This confirmed that novelty was similarly valued in each valence context. Across sessions, the model-derived choice probabilities were positively correlated with the monkeys’ actual choice behavior (average *r* = .59; *t*(106) = 53.18, *p* < .001, 95% CI [.56, .61]; **Figure 3D**), especially when model predictions were averaged to reflect behavioral assessments of the monkeys’ choice behavior (**Figure 3A-C**).

**Figure 3.**
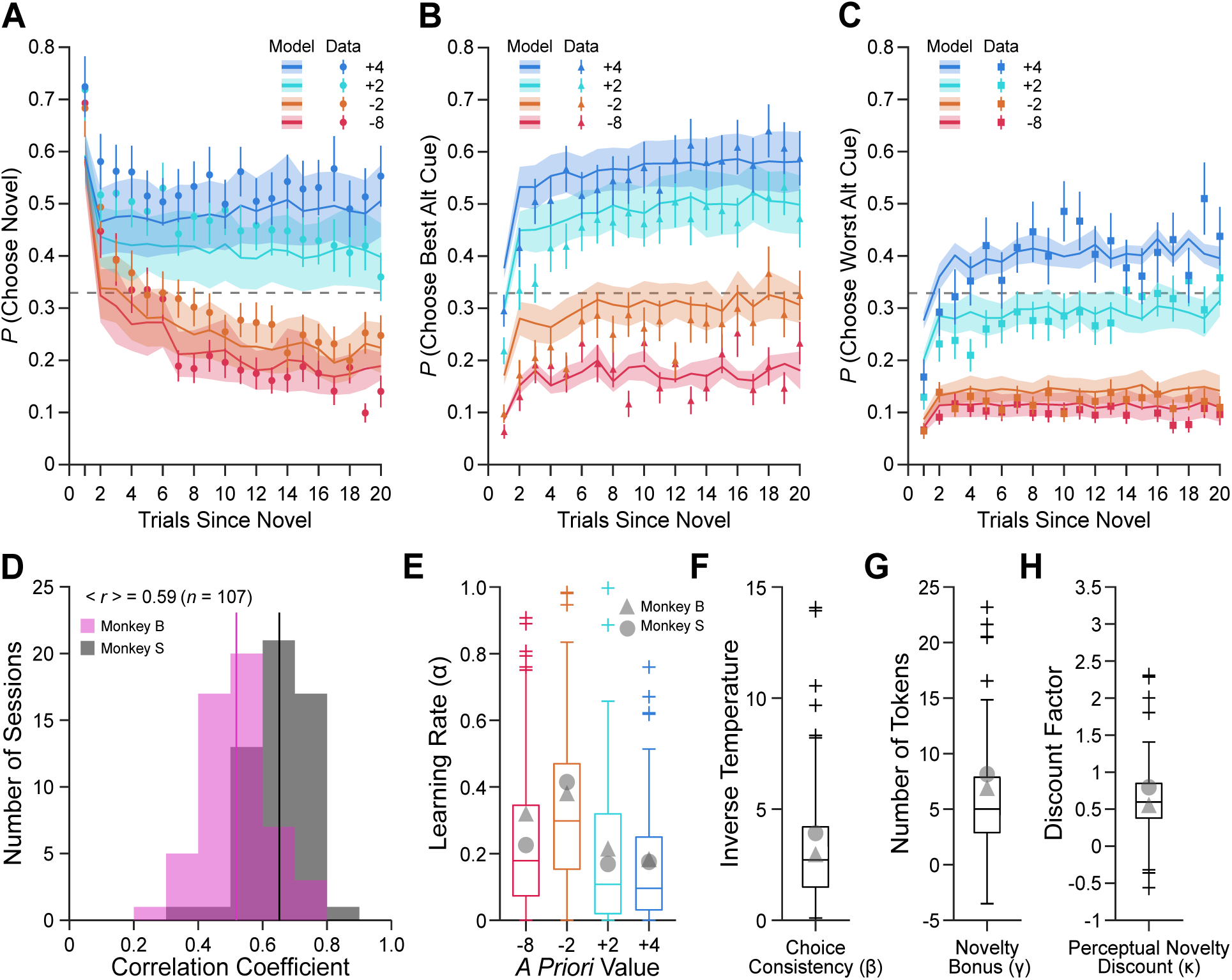
Reinforcement learning model with novelty bonus revealed monkeys learned faster from losses compared to gains. **(A)** Solid lines indicate the model estimates of the probability of choosing the novel option as a function of trials since the novel cue was introduced, separated by the *a priori* value of the novel cue. Shaded area is the S.E.M. of the model estimates across sessions. Markers show the behavioral probability of choosing the novel option and error bars are the S.E.M. across sessions. Grey dashed line is 33%, or chance choice probability across the three available cues, for panels **A**, **B** and **C**. **(B)** Solid lines indicate the model estimates of the probability of choosing the best alternative option as a function of trials since the novel cue was introduced, separated by the *a priori* value of the best alternative cue. Shaded area is the S.E.M. of the model estimates across sessions. Markers show the behavioral probability of choosing the best alternative option and error bars are the S.E.M. across sessions. **(C)** Solid lines indicate the model estimates of the probability of choosing the worst alternative option as a function of trials since the novel cue was introduced, separated by the *a priori* value of the worst alternative cue. Shaded area is the S.E.M. of the model estimates across sessions. Markers show the behavioral probability of choosing the worst alternative option and error bars are the S.E.M. across sessions. **(D)** Histogram of session-by-session correlation coefficients for the behavioral data probability of choosing the novel, best alternative, and worst alternative option across the first 20 trials since the novel option was introduced and their matched model estimates. The pink histogram represents Monkey B’s correlation coefficients (*n* = 53 sessions), and the grey histogram represents Monkey S’s correlation coefficients (*n* = 54 sessions). The solid lines represent each monkey’s average correlation coefficient across sessions. The average correlation coefficient was .59. **(E)** Best fitting model learning rates (α) separated by *a priori* cue value. Box plots represent session estimates from both monkeys; gray markers indicate individual monkey averages. **(F)** Best fitting model choice consistency values (β). Box plot represents session estimates from both monkeys; gray markers indicate individual monkey averages. **(G)** Best fitting model novelty bonus values (γ). Box plot represents session estimates from both monkeys; gray markers indicate individual monkey averages. **(H)** Best fitting model perceptual novelty discount (κ). Box plot represents session estimates from both monkeys; gray markers indicate individual monkey averages.

The model also estimated four learning rates (**Figure 3E**) based on the four possible value categories to which a cue was assigned (Taswell et al., 2018). Direct comparisons of the estimated learning rates indicated that the monkeys learned faster from loss compared to gain cues. We found that overall, cues associated with losses had higher learning rates than cues associated with gains (**Figure 3E**; Valence: *F*(1, 105) = 16.59, *p* < .001). However, it was clear that this effect was driven by heightened learning rates for cues associated with small losses. Learning rates were higher for cues associated with small losses compared to large losses (−2 vs. −8, *t*(105) = 3.038, *p* = .003) and small (−2 vs. +2, *t*(105) = 4.43, *p* < .001) or large gains (−2 vs. +4, *t*(105) = 4.98, *p* < .001). Whereas the learning rates for the two categories of gain cues did not differ (+2 vs. +4, *t*(105) = 0.45, *p* = .66). These results suggest the monkeys were motivated to avoid losing tokens, consistent with prior evidence that macaques’ decision making aligns with prospect theory (Kahneman & Tversky, 1979; Yang et al., 2022).

### Task Engagement Reflects Shifts in Overall Motivation

The monkeys’ avoidance of cues associated with small or large losses suggests that losing tokens was aversive, influencing their motivational state. Changes in motivational state should have broader effects on task performance. Sequential or frequent losses in a block of trials made it difficult to accrue and cash out tokens for primary rewards, and this should reduce the monkeys’ motivation to engage with the task. A refusal to initiate or hold central fixation is one measure used to track changes in macaques’ motivational state (Burk et al., 2024; Taswell et al., 2023).

To examine what triggered central fixation aborts, we tabulated the events that preceded them in our task. We found that there were five main events that preceded the monkeys aborting central fixation (**Figure 4A**). By far, the most frequent event preceding a central fixation abort was a cash out of the accumulated tokens (Monkey B: 52%, Monkey S: 39%; *t*(106) = 18.43, *p* < .001). The next two most frequent events were a loss of tokens (B = 19% / S = 17%) and a gain of tokens (B = 16% / S = 20%), both of which were less frequent than chance (lost tokens: *t*(106) = −12.67, *p* < .001; gained tokens: *t*(106) = −12.57, *p* < .001) and occurred with equal frequency to one another (*t*(212) = 0.098, *p* = .92). The fourth most frequent event preceding a central fixation abort was receiving no feedback on the previous trial due to starting the trial with 0 tokens and choosing a loss cue (B = 8% / S = 12%; *t*(106) = −21.74, *p* < .001). Finally, central fixation aborts also occurred on trials in which the monkeys accumulated more than 12 tokens, but the tokens were not cashed out, because the preceding trial included a cash out (Monkey B = 5%; Monkey S = 13%; *t*(106) = −25.13, *p* < .001). These events likely reflect a decrease in the value of engaging with the task due to motivational shifts (Burk et al., 2024).

**Figure 4.**
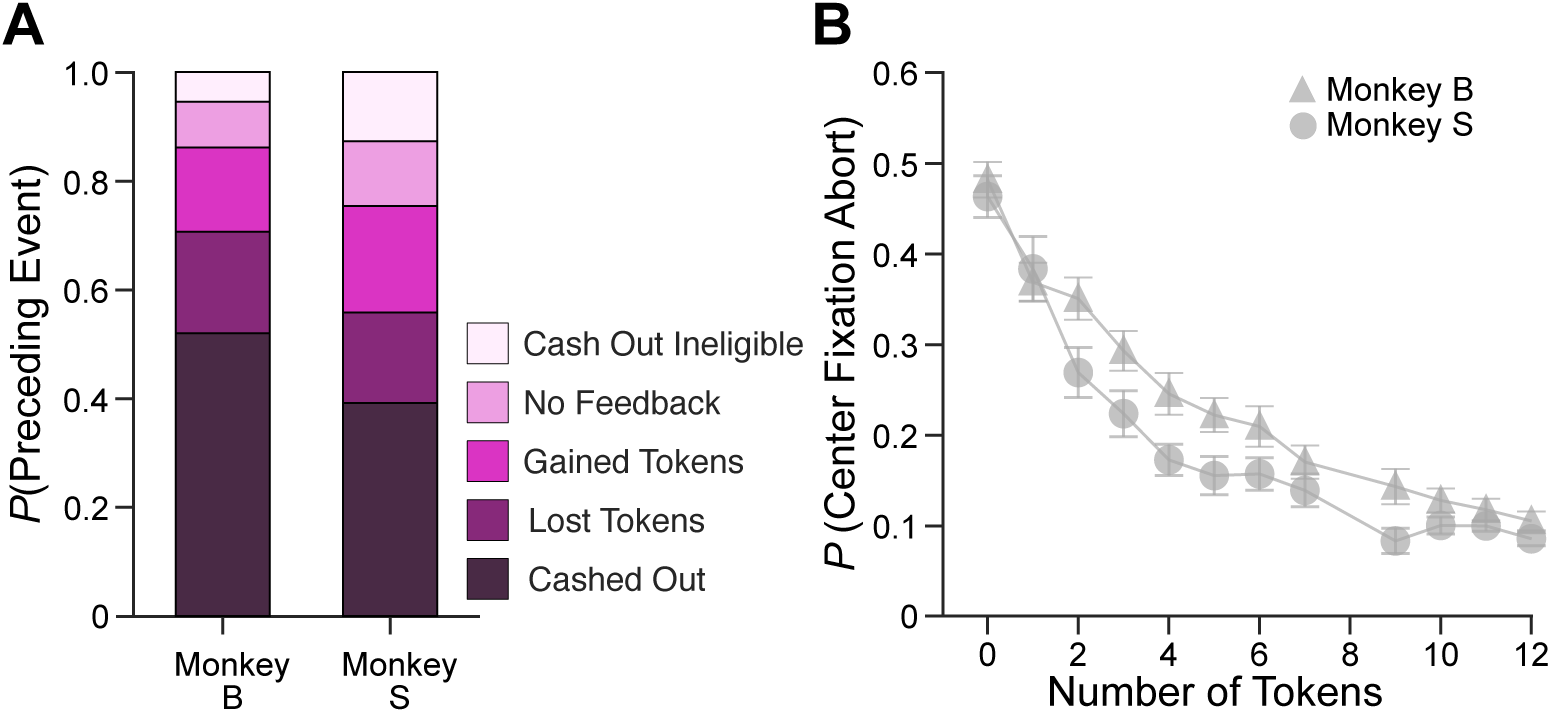
Center fixation aborts illustrate motivational relevance of tokens. **(A)** Probability that a center fixation abort was preceded by five different events, separated by monkey. **(B)** Probability of aborting following the initiation of central fixation as a function of number of tokens at the start of the trial.

A prior study has found that central fixation aborts are negatively correlated with the number of tokens accumulated at trial start (Burk et al., 2024). Collapsing across all the preceding event types, we also found a negative correlation between central fixation abort frequency and the number of tokens accumulated at the start of the trial (**Figure 4B**; average *r* = −.57; *F*(1, 107) = 508.25, *p* < .001). The monkeys aborted central fixations most often when they started a trial with 0 tokens and this behavior declined exponentially as they accumulated more tokens and there was an increased probability of cash out. So, while the monkeys’ motivation did not influence novelty seeking, it clearly affected task engagement.

### Choice Balks Reflect Strategic Responses to Approach-Avoidance Conflicts and Uncertainty

Besides central fixation aborts, we also observed that the monkeys would frequently balk their choices after fixating a particular choice option (~20% of trials; **Figure 5**). We initially investigated if choice balks were another motivational indicator. But unlike central fixation aborts, how often the monkeys balked their choices did not vary with the number of tokens accumulated at the start of the trial (**Figure 5A**; *F*(1, 108) = 1.82, *p* = .18). To better understand why the monkeys balked their choices, we examined the *a priori* value of the choices that elicited the balks. Balks were more frequent when the monkeys had selected an option associated with a loss than a gain (**Figure 5B**; *F*(3, 315) = 670.69, *p* < .001). Moreover, within each valence category balks were more or less frequent based on the magnitude of the token loss (large vs. small loss: *t*(105) = 8.65, *p* < .001) or gain (large vs. small gain: *t*(105) = −3.92, *p* < .001). This suggests balks were deliberate attempts to optimize outcomes by avoiding anticipated losses or selecting more favorable alternatives.

**Figure 5.**
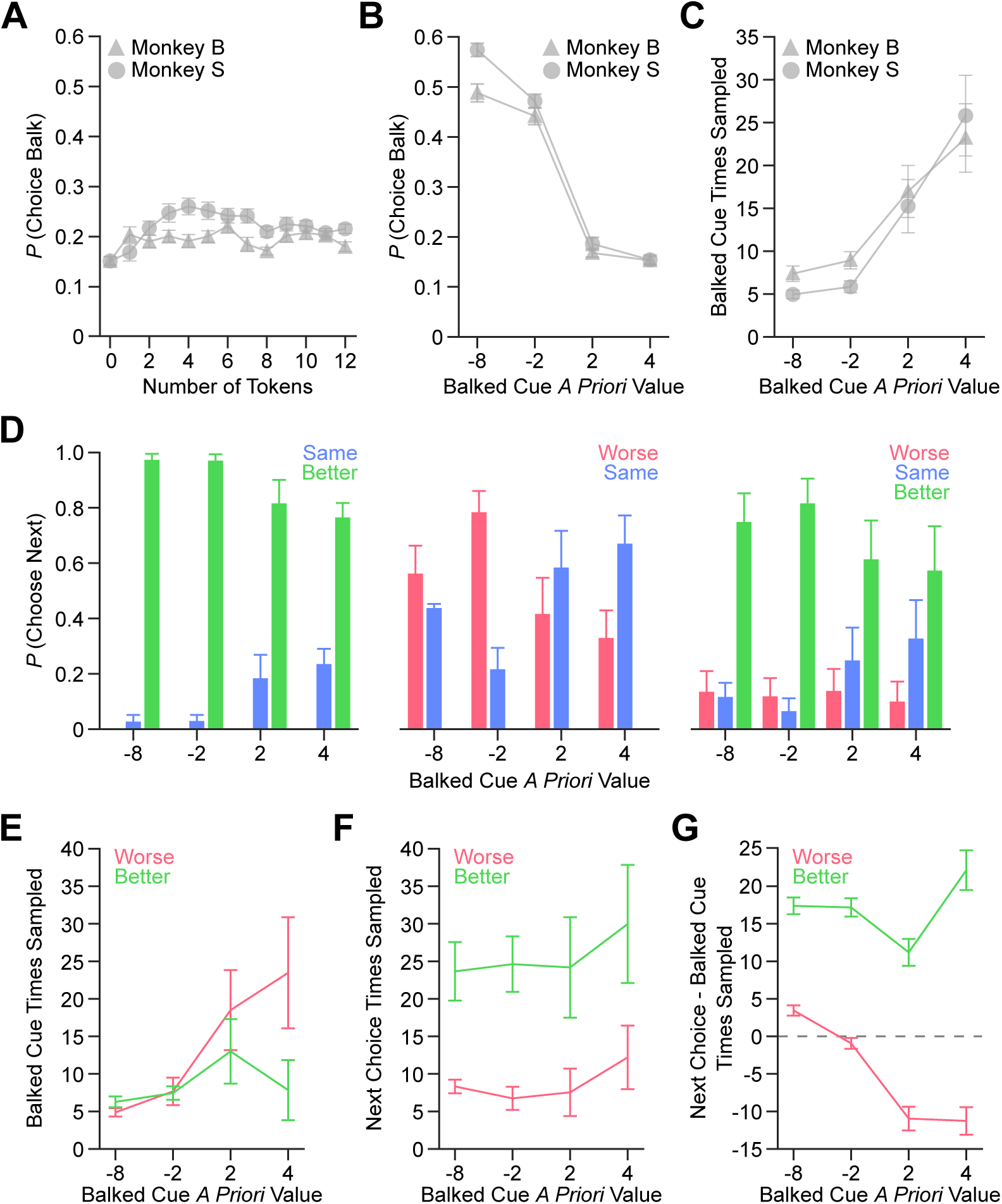
Choice balks and subsequent choices show valence-dependent exploratory strategies. **(A)** Probability of balking fixation on a cue after the monkey had begun fixating on it (choice balk), by the number of tokens at the start of the trial, separated by monkey. Error bars are S.E.M across sessions. **(B)** Probability of balking on a choice by the value of the cue the monkey had begun fixating on, separated by monkey. Error bars are S.E.M across sessions. **(C)** The average times the cue balked on had been sampled by the value of the balked cue, separated by monkey. Error bars are S.E.M across sessions. **(D)** Probability of choosing a cue that was worse (lower value, pink), the same (repeat choice, blue), or better (higher value, green) on the next completed trial following a choice balk, separated by the value of the cue that they balked on. Each panel represents a different condition they balked in: (left) they could choose one of two better options or repeat the choice they balked on, (center) they could choose one of two worse options or repeat the choice they balked on, (right) they could choose a worse option, repeat their choice, or choose a better option. Error bars are S.E.M across sessions. **(E)** Average number of times the cue balked on was sampled prior to the balked choice, separated by the value of the balked cue and the next correct choice’s value relative to the balked cue. Error bars are S.E.M across sessions. **(F)** Average number of times the next chosen cue following a choice balk was sampled, separated by the value of the balked cue and the next correct choice’s value relative to the balked cue. Error bars are S.E.M across sessions. **(G)** The subsequent choice number of times sampled following a balk minus the balked cue number of times sampled as a function of the balked cues’ average value, separated by the subsequent correct choice being worse or better than the cue they balked on. Values above zero signify the subsequent choice had been more frequently sampled than the balked cue and was less informative. Values below zero signify the subsequent choice had been less frequently sampled than the balked cue and was more informative. Values around zero signify the subsequent choice had been equally sampled compared to the balked cue and was not more or less informative. Error bars are S.E.M across sessions.

To test this hypothesis, we reexamined the balks in terms how many times the monkeys had sampled an option, prior to its selection eliciting a balk. Balked cues associated with a large loss were sampled less than balked cues associated with a small loss (**Figure 5C**; *F*(1, 105) = 14.31, *p* < .001), consistent with the monkeys attempting to limit further selection of loss cues based on their *a priori* value. Balked cues associated with a large gain were sampled more than balked cues associated with a small gain (*F*(1, 105) = 56.9, *p* < .001). However, the balked selection of cues associated with larger gains was puzzling.

Balking a choice was not advantageous from the standpoint of maximizing rewards over time. This is because the monkeys had to repeat the trial. It would be advantageous, however, if the monkeys were attempting to maximize gains or minimize losses to increase their probability of having the tokens cashed out over a short time horizon. To further contextualize the monkeys’ balking behavior we examined whether the monkeys chose a cue whose empirical value was worse, the same, or better than the empirical value of the balked choice. Whether a better or worse choice option was available depended on the choice environment (**Figure 5D**). When the monkeys’ balked choice was the least valuable option available (*n* = 5,882 trials), the monkeys had to next choose either the same or a better option (**Figure 5D** left panel), and the monkeys frequently chose to select the better option. But as the *a priori* value of the balked cue increased, the monkeys were less likely to choose a higher value option (*F*(3, 233) = 18.68, *p* < .001) because they were more likely to repeat their choice (*F*(3, 184) = 42.59, *p* < .001). Overall, these balks could stem from lapses in attention or response location biases.

When the monkeys’ balked choice was the most valuable option available (**Figure 5D** center panel; *n* = 3,209 trials), the monkeys had to next choose either the same or a worse option. In these instances, we observed a valence-dependent pattern in the monkeys’ willingness to explore a worse option. When the monkeys balked a choice associated with a loss, their next choice was more likely to be an option lower in value than the cue that had elicited the balk. Whereas when they balked a choice associated with a gain, they were more likely to repeat their choice than to choose a worse option (*F*(3, 184) = 42.59, *p* < .001). It is more difficult to ascertain what causes these types of balks.

When the monkeys’ balked choice was neither the best or worst option available to them, the monkeys had greater flexibility in what they chose next (**Figure 5D** right panel; *n* = 3,640 trials). The monkeys could either repeat the balked choice or choose an option higher or lower in value. The monkeys were more prone to choose a better option than a worse option (*F*(1, 110) = 985.39, *p* < .001) and did so more often when the balked choice was associated with losing tokens (*F*(3, 284) = 22.48, *p* < .001). Likewise, they were more likely to repeat their choice when they balked selection of a gain cue (*F*(3, 284) = 33.35, *p* < .001). But the *a priori* value of the balked choice did not influence how often the monkeys subsequently selected a worse option (*F*(3, 284) = 0.98, *p* = .41).These balks are likely intended to improve the trial outcome over what could have occurred if the choice was maintained.

Balking their choices to select lower value options (**Figure 5D** middle panel) contradicts the hypothesis that the monkeys were simply seeking a second opportunity to maximize gains and minimize losses (i.e., resolving approach-avoidance conflicts). We hypothesized that balking a choice could additionally be related to uncertainty about option values. This could explain why the monkeys would select lower value options (i.e., explore) rather than the same or higher value options (i.e., exploit). Uncertainty about choice outcomes might also explain why the monkeys chose to balk choices associated with large gains.

To test this hypothesis, we reexamined the mean number of times a balked option was sampled prior to the balk occurring (**Figure 5C**). We did so based on whether monkeys next chose an option that was higher or lower in value (**Figure 5E**). When the monkeys balked choices associated with gaining tokens, the value of the option they chose next was related to how many times they had sampled the balked choice option. Whereas when they balked choices associated with losing tokens, what they chose next was unrelated to sampling of the balked choice. This suggests that uncertainty about choice outcomes only influenced what monkeys chose next when they balked selection of a gain cue. Overall, the number of times they had previously sampled the option they chose after balking was greater when they chose a better compared to a worse option (**Figure 5F**).

To determine if the monkeys balked their choices to choose a more or less informative option, we subtracted the number of times they had sampled the option that elicited the balk from the number of times they had selected the option chosen after a balk. We examined this difference based on whether the option chosen after a balked choice was higher (better) or lower (worse) in value (**Figure 5G**). When the monkeys balked their choice and selected a better option, it was to exploit an option whose outcome was more certain. Whereas, when the monkeys balked choices and selected a worse option, it was to gain information about cues they had sampled less frequently and whose value was less certain (i.e., exploration). We found that exploratory choices following balks were valence dependent, while exploitative choices were not (**Figure 5G**; *F*(3, 215) = 22.34, *p* < .001). When the monkeys balked choices associated with gains and subsequently chose a worse option, the worse option was sampled less than the choice option that elicited the balk (*t*(202) = −12.38, *p* < .001). While this could lead to a worse outcome, it ensured a gain in information that could be exploited in the future. Whereas, when the monkeys balked choices associated with losses and subsequently chose a worse option, the worse option was sampled no more or less than the option that elicited the balk (*t*(136) = 0.88, *p* = .38).

These findings highlight the strategic nature of the monkeys’ decisions, which reflect both reward maximization and uncertainty reduction. At a superficial level, these balks are related to resolution of approach-avoidance conflicts intended to maximize reward. However, a closer examination demonstrates they are also a useful strategy to gain information and solve the explore-exploit dilemma.

## Discussion

We demonstrate that macaques learned to efficiently manage explore-exploit tradeoffs when exploration was associated with both appetitive (token gains) and aversive (token losses) outcomes. Monkeys did explore novel, uncertain opportunities more often when exploitation of previously learned values would result in losing rather than gaining tokens. But this difference was due to an optimistic valuation of novelty and not a valence-dependent shift in the monkeys’ overall willingness to explore. While hedonic valence did not modulate exploration, it did modulate reinforcement learning. The monkeys were quicker in learning to avoid choosing options that resulted in token losses compared to token gains. The valence of the decision environment also influenced the monkeys’ motivation and elicited strategic decisions to balk choices when there was potential to make a more informed choice that minimized token losses.

### Influence of Novelty Value and Outcome Valence on Directed Exploration of Novelty

The monkeys’ novelty-driven exploration is a form of directed exploration (Costa et al., 2019; Wilson et al., 2021), since novel options, when they are first introduced, are always sampled less than familiar options whose value is already learned. The prevailing view, based on human research using verbal instructions about the goal of exploring, is that directed exploration is heightened in loss contexts (Blanchard & Gershman, 2018; Krueger et al., 2017; Wilson et al., 2014). While we did find evidence for this view (**Figure 2C**), it was clear in modeling the monkeys’ choices to explore or exploit that they deployed a unified value-based strategy for exploration, independent of the valence context (**Figure 2E**). Furthermore, that they were relying on an optimistic prior belief that resolving uncertainty is valuable (**Figure 3G**). Moving forward, it will be critical to understand how exploration is influenced by motivational priors formed either through verbal instructions and or experience. For instance, dopamine manipulations in monkeys enhance novelty valuation after experience with rewarding outcomes (Costa et al., 2014), whereas similar manipulations in humans reduce directed exploration (Chakroun et al., 2020) or blunt the influence of valence on information-seeking (Vellani et al., 2020). Future studies should explicitly manipulate prior beliefs about novelty value to better characterize how biases in novelty and information seeking lead to imbalances in explore-exploit decision making.

The pattern of the monkeys’ novelty-driven exploration also suggests that novelty seeking is predicated on gaining information to resolve uncertainty about the decision environment, and that novelty seeking occurs even in aversive contexts. Monkeys and humans are especially novelty prone throughout their lifespan (Bliss-Moreau & Baxter, 2019), and continued use of directed exploration is a sign of healthy cognitive aging (Mizell et al., 2024). But other species are less neophilic, and many are even neophobic. One advantage to the use of multi-armed bandit tasks where novel options are introduced to induce explore-exploit tradeoffs is that the tasks are easily implemented across species (Hogeveen, Mullins, et al., 2022). Cross-species comparisons of valence asymmetries in novelty-seeking behavior could determine if the optimistic valuation of novelty is specific to primates (Krueger et al., 2017; Leopold & Averbeck, 2022). This would aid in identifying the potential neural bases of Bayesian shrinkage.

### Influence of Valence on Directed and Random Exploration to Resolve Motivational Conflicts

Besides novelty-driven exploration, we also observed that monkeys made decisions to explore or exploit after they balked a choice. This required them to repeat the trial with knowledge of the choices available to them, but without experiencing a choice outcome. When the monkeys balked their choice and selected a better option, it was to exploit an option whose outcome was more certain. Whereas, when the monkeys balked choices and selected a worse option, it was to gain information about cues they had sampled less frequently and whose value was less certain. These self-generated exploratory bouts are seemingly triggered by approach-avoidance conflicts. This has not been observed in prior examinations of macaque explore-exploit decision making in exclusively appetitive contexts and involving primary rewards (Costa et al., 2014, 2019; Ebitz et al., 2018; Giarrocco et al., 2024). Although exploration does occur independently from exogenous changes in the decision context, such as in drifting multi-armed bandit tasks (Daw et al., 2006; Ebitz et al., 2018; Speekenbrink & Konstantinidis, 2015). Exploratory choices following choice balks resemble directed exploration. The frequency of choice balks that triggered the exploratory events were not linked to the introduction of a novel option or the number of tokens accrued at the start of the trial, implying tonic use of an information-seeking strategy (Ebitz et al., 2018). But directed exploration was only observed following the monkeys balking choices associated with gains. The subsequent choices following the monkeys balking choices associated with losses did not provide more information and therefore represented random exploration.

### Potential Insights into Neural Mechanisms of Explore-Exploit Decision Making

The neural mechanisms underlying directed and random exploration are increasingly found to involve a complex interplay between prefrontal cortical, and subcortical motivational neural circuits. The dorsolateral prefrontal cortex and frontopolar cortex have been implicated in mediating exploratory decision-making in humans (Daw et al., 2006; Ebitz et al., 2018; Zajkowski et al., 2017), particularly directed exploration (Zajkowski et al., 2017). Neurons in macaque orbitofrontal cortex, amygdala, and ventral striatum also encode value signals that predict the use of directed exploration (Costa & Averbeck, 2020; Costa et al., 2019), and these findings were replicated in humans using neuroimaging (Hogeveen, Mullins, et al., 2022). Moreover, decisions to explore or exploit are encoded in motivational circuits after a choice is made and feedback received, whereas the same decisions are encoded in dorsolateral prefrontal cortex prior to a choice being made (Tang et al., 2022). The monkeys’ consistent use of directed exploration across valence contexts suggests that prefrontal circuits provide a general exploration signal that is subsequently modulated by encoding of hedonic valence and motivational signals in corticolimbic circuitry.

While novelty-driven directed exploration was not valence-dependent, self-generated bouts of directed exploration after balking a choice were more prevalent in appetitive contexts (**Figure 5G**). Whereas self-generated bouts of random exploration after choice balks were specific to aversive contexts. The frontopolar cortex is routinely implicated in mediating directed exploration and in evaluating feedback from self-generated decisions (Tsujimoto et al., 2010). Previous work has also shown that random exploration is not attributable to the frontopolar cortex (Zajkowski et al., 2017), and is instead due to encoding of decision noise in dorsolateral prefrontal cortex (Ebitz et al., 2018). This raises additional questions about whether distinct appetitive and aversive neural circuits are specifically engaged by prefrontal cortical circuitry mediating explore-exploit decision making in particular contexts.

### Approach and Avoidance Motivation Influence Reinforcement Learning

Secondary reinforcers offer a powerful tool for manipulating motivational valence in decision-making tasks, allowing gain and loss outcomes to be precisely equated. In our task, monkeys explored novel options but also learned to avoid cues that resulted in token losses more quickly than they learned to favor cues that resulted in token gains. This valence asymmetry in reinforcement learning (**Figure 3E**) fits with evidence of asymmetries in learning and motivation using primary reinforcers in rodents (Beyeler et al., 2016; Miller, 1959; Namburi et al., 2015). Previous studies using primary reinforcers in nonhuman primates (e.g., food, juice, or airpuff) have struggled to equate appetitive and aversive outcomes, limiting the ability to probe how valence shapes learning. By contrast, secondary reinforcers (e.g., virtual tokens) enable the computation of losses and gains in a common motivational currency. Prior studies that used tokens to examine rhesus macaques’ learning and decision making in multi-armed bandit tasks where token cash out was randomized, did not find that monkeys discriminated between cues associated with different magnitude losses (Burk et al., 2024; Taswell et al., 2018). In our task, losses were more salient than in prior studies, because the monkeys needed to avoid losses to accumulate tokens in order to increase the probability of exchanging their tokens for juice.

Because the monkeys’ choice behavior indicated that losing tokens was aversive, we looked for additional signs of changes in their motivational state outside of choice behavior. We considered central fixation aborts as a potential proxy. In nonhuman primates, trial aborts are more frequent when effort is high (Pasquereau & Turner, 2013; Varazzani et al., 2015), or reward probability is low (Burk et al., 2024; Inaba et al., 2013; La Camera & Richmond, 2008). Here, losing tokens reduced the likelihood of exchanging those tokens for juice, increasing the effort and delay to reach the cash out threshold (**Figure 1D**). Not surprisingly, then, central fixation aborts were highest when the monkeys started a trial with no tokens and declined as they accrued tokens (**Figure 4**). The monkeys’ increased tendency to balk choices when they anticipated losing compared to gaining tokens further suggested they were trying to avoid losing tokens, and entering an extended, aversive motivational state (Averbeck & Murray, 2020; Bliss-Moreau & Rudebeck, 2021; Burk et al., 2024; Eldar et al., 2016).

Together, these findings show that monkeys’ explore-exploit decision-making is shaped by approach-avoidance conflicts, with heightened sensitivity to loss influencing both learning and task engagement. The ability to use secondary reinforcers to control hedonic valence and arousal within a reinforcement learning framework opens new opportunities for studying motivational circuits across species using translationally relevant tasks (Hogeveen, Mullins, et al., 2022).

### Constraints on Generality

These findings are based on a study of two adult male rhesus macaques performing a token-based reinforcement learning task. As such, the generality of our conclusions is subject to several important constraints. First, the use of a secondary reinforcer may not fully capture the motivational salience of primary rewards or punishers, but this same issue exists for many human studies that motivate participants with gaining or losing points or monetary rewards. So, it remains unclear whether similar patterns of exploration and learning would emerge under different reinforcement modalities and when the same processes are studied in different species. However, monkeys exhibit similar patterns of novelty-driven exploration when they complete multi-armed bandit tasks for primary rewards (Costa & Averbeck, 2020; Costa et al., 2019) and their behavior aligns with novelty-driven exploration exhibited by humans completing the same task for secondary reinforcers (Hogeveen, Mullins, et al., 2022). Second, the study included only male subjects from a single primate species, so the observed strategies may not extend to females or to species with differing cognitive or affective capacities. While there is evidence of sex differences in explore-exploit decision making in mice (Chen et al., 2021), higher cognitive function does not differ between male and female rhesus monkeys using tests of episodic memory and strategy implementation (Nagy et al., 2017). Third, we used the novel choice options to introduce non-stationarity into the choice environment, and determining if exploration triggered by other forms of uncertainty (e.g., volatility or ambiguity) requires further investigation.

## Conclusion

These results illustrate that monkeys use directed and random exploration in the same way humans do, to support learning and adaptive decision making in uncertain environments. They also demonstrate that monkeys efficiently solve the explore-exploit dilemma while managing approach-avoidance conflicts that arise when choosing between options associated with different valence outcomes. An integrative understanding of motivated decision making in the domains of gains and losses can illuminate how these processes can become maladaptive in anxiety and mood disorders.

## Methods

### Subjects

Two male rhesus macaques (*Macaca mulatta*) were studied (Monkey S: 6.2 years old, 9.5 kg; Monkey B: 6.7 years old, 10.4 kg). All monkeys were placed on water control for the duration of the study and on test days earned all their fluid through performance on the task. Experimental procedures for all monkeys were performed in accordance with the National Institutes of Health Guide for the Care and Use of Laboratory Animals and were approved by the Oregon National Primate Research Center (ONPRC) Animal Care and Use Committee.

### Experimental Setup

All monkeys’ actions and choices were controlled through operant conditioning and monitoring of their eye movements. Stimuli were presented on a 19-inch LCD monitor situated 40 cm from the monkeys’ eyes. During training and testing, each monkey was seated in a primate chair with their heads restrained. Stimulus presentation and behavioral monitoring were controlled by a PC computer running Monkeylogic (Asaad & Eskandar, 2008), a MATLAB-based behavioral control program. Eye movements were monitored at 400 fps using an Arrington Viewpoint eye tracker (Arrington Research, Scottsdale, AZ) and sampled at 1 kHz. A gravity-fed reward system controlled by a computer-controlled solenoid valve (Mitz, 2005) delivered a fixed amount of reward (0.09 mL on average per solenoid opening, 95% CI: 0.06-0.15mL/opening) as described below.

### Secondary Reinforcement Training

Monkeys were first trained on a deterministic two-arm bandit task in which choices were associated with differently sized primary juice rewards (Costa et al., 2014, 2019). Once animals consistently reached criterion (~80% accuracy in selecting the higher value option), we added tokens as secondary reinforcers that would be given following their choices, then immediately cashed out at the end of the trial. Once the monkeys reached criterion in this task iteration, we introduced different size token gains to facilitate them in learning based on value comparisons. After reaching criterion in this training iteration, we required that the monkeys completed two trials before cash out, then three, then four, then five. After stepping through all the fixed ratio schedules up to five trials and reaching criterion, we introduced an endowment of eight tokens for the monkeys. We then introduced cues that could result in a loss in their token endowment and used the same procedure for learning different sized gains for the loss cues. Once they had reached criterion in learning between differently sized losses, we moved them into the final version of the three-arm bandit task, described below.

### Task Design and Stimuli

The monkeys were trained to complete a three-arm bandit task to acquire tokens that were cashed out for juice reward (**Figure 1**). The monkeys were trained prior to the task to associate the exchange of individual tokens with receipt of a primary reward, a fixed amount of apple juice (0.09mL/token). The task design was an adaption of a multi-armed bandit task previously used in humans and monkeys to study explore-exploit trade-offs (Costa et al., 2014, 2019; Djamshidian et al., 2011; Hogeveen, Mullins, et al., 2022; Wittmann et al., 2008). The main difference is that the monkeys were first trained to regard visual tokens as secondary reinforcers that could be gained or lost based on their performance in the task.

Before each trial, three images were randomly drawn from a set of four images. Randomized trial-by-trial selection of the images was intended to limit the monkeys’ ability to predict the choice options and delay their decision making until the choice cues were presented. The trial began with displaying the current token endowment at the bottom of the computer display, followed by the presentation of a central fixation stimulus. On each trial, the monkey had to acquire and hold central fixation for a variable length of time (500-750 ms). Failing to acquire or hold central fixation aborted the trial and accrued a 1 s delay until the trial restarted (i.e., central fixation aborts). Aborting central fixation did not allow the monkeys to advance to the next trial. After holding central fixation, three peripheral choice targets were presented at the vertices of an upward facing triangle. The locations of the three stimuli were randomized from trial-to-trial. The monkeys’ first saccade away from central fixation stimulus to one of the three choice targets was registered as their choice. To confirm their choice the monkeys were required to maintain fixation of the chosen target for 500 ms. A saccade away from the chosen target before 500 ms elapsed aborted the trial and accrued a 1 s delay until the trial restarted (i.e., choice balks). The location and identity of the choice options did not change when the trial restarted. Balking a choice therefore did not allow the monkeys to advance to the next trial.

Each image was randomly assigned a normal distribution of token outcomes with a different mean but the same standard deviation (SD = 1). The *a priori* value of each image was based on the mean of this distribution and took on one of four values: −8, −2, +2, and +4. The distributions were truncated such that cues associated with gains (+2 and +4) never yielded a loss of tokens and the cues associated with losses (−8 and −2) never yielded a gain of tokens. The number of tokens earned after choosing a gain or loss cue was based on a discrete token value drawn from the associated distribution.

The monkeys started each session with an endowment of 8 tokens. From this point, they either gained or lost tokens from one trial to the next based on their choices. The number of tokens at the end of trial determined the probability that all the displayed tokens would be automatically cashed out and a corresponding amount of juice dispensed. When the monkeys had 4 tokens or fewer, the probability of having the tokens cashed out equaled 0. When the monkeys had accumulated between 5 and 11 tokens, the probability of having the tokens cashed out increased linearly. This probability was based on the integration of a discrete uniform distribution, U {5,12}. When the monkeys had accumulated 12 or more tokens the probability of having the tokens cashed out equaled 1. Disappearance of each token coincided with opening of the solenoid valve and delivery of a fixed amount of juice reward. After cash out, the token endowment was replenished with 8 tokens, and this was indicated to the monkey by displaying the tokens at the bottom of the screen on the next trial.

The monkeys did not incur a debt. If a monkey started a trial having accumulated 0 tokens and chose a loss cue, it started the next trial with 0 tokens. Likewise, if they started a trial with fewer tokens than they lost following a choice, they started the next trial with 0 tokens.

When a novel stimulus was introduced, it replaced one of the existing choice options. Which alternative option it replaced was randomly determined, with the constraint that no single choice option could be available for more than 160 consecutive trials. The trial interval between the introduction of two novel stimuli followed a discrete uniform distribution, U {10,30}. Novel choice options were randomly assigned one of the four reward distributions. The only constraint was that one out of the four options available after introducing a novel option was a gain cue. The monkeys were more likely to experience trials with the opportunity to gain rather than lose tokens (ANOVA; Trial Type: *F*(1,320) = 9.57, *p* < .01). Each trial contained three options, and we categorized these based on whether two or more cues would result in losses, or two or more cues would result in gains. Specifically, 56.3% of trials provided more opportunity for gaining than losing tokens (Loss/Gain/Gain and Gain/Gain/Gain), while the remaining 43.7% of trials had a larger probability for losing tokens (Loss/Loss/Loss and Loss/Loss/Gain). Additionally, pure valence trials, or the trials with all loss or all gain cues, were relatively rare (5.4% and 11.5%, respectively). However, to maintain monkeys’ motivation, we limited the number of consecutive Loss/Loss/Loss trials to 8. As such, there were more trials consisting entirely of gain compared to loss cues (*F*(1,106) = 41.65, *p* < .001). During the more frequently encountered mixed valence trial types, where the monkeys are provided with a combination of gain and loss cues, they were more likely to encounter Loss/Gain/Gain trials (44.8%) than Loss/Loss/Gain trials (38.3%) (*F*(1,105) = 16.9, *p* < .001).

Each daily session utilized a unique set of 48 pictures, randomly selected from the THINGS database (Hebart et al., 2019). The THINGS database is publicly available at https://things-initiative.org. The images were screened for image quality, discriminability, uniqueness, size, and color and never repeated across sessions. The first block of trials in each session utilized four randomly selected images, after which a novel image was pseudorandomly introduced, forming a new block and set of four images. There were four images per block, four novel images in the first block and one new image for every following block, and up to 45 blocks per session. Images were never repeated across sessions. We collected 53 behavioral sessions from Monkey B and 54 from Monkey S, with an average of 28 blocks per session (Monkey B: 30; Monkey S: 26) and an average of 540 completed trials per session (Monkey B: 580; Monkey S: 499).

### Strategy for Statistical Analyses

We quantified choice behavior as the fraction of times the monkeys chose either the novel or the best alternative option as a function of how many trials since the novel option was introduced – separated by the *a priori* value of the cues. The best alternative option was not fixed trial-to-trial and instead, was defined as whichever of the known alternative options in a trial with the novel cue was the highest expected value. In cases where the remaining options were equal in their *a priori* values (this occurred 19% of the time), the best alternative option was defined as the cue with the highest empirical value between the two. We exclude the first block of every session from analyses, as all four of the cues presented during the first block are novel to the monkey.

For statistical analyses, each dependent variable was entered into a full factorial, mixed effects ANOVA model implemented in MATLAB or SAS JMP. For all ANOVA models, monkey identity and testing session were random effects, with testing session hierarchically nested under monkey identity. For the analyses of the distribution of valence combinations in trials, the model included fixed effects of trial type (LLL/LLG/LGG/GGG, L: loss, G: gain). For the analyses of choice behavior, the model included fixed effects of *a priori* cue value (−8, −2, +2 +4) or valence, and some models included trials since the novel option was introduced. For the analyses on learning rates from the reinforcement learning model, we used valence or *a priori* cue value as fixed effects. For analyses of central fixation aborts, the model included a fixed effect of number of tokens at the start of the trial, which was specified as a continuous variable. For the analyses of choice balks, the model could include fixed effects of number of tokens at the start of the trial, *a priori* cue value or the valence of the option they balked on, whether they chose a worse or better option on the next trial, and the number of times the balked cue or the subsequently chosen cue had been sampled. Post hoc analyses of significant main effects used Fisher’s least significant difference test to correct for multiple comparisons (Levin et al., 1994). Post hoc tests of significant interactions and pairwise comparisons required computing univariate ANOVAs for individual component effects and correcting for multiple comparisons.

### Reinforcement Learning Model

Previous work using multiple values across valences in a bandit task led to our choice of fitting four learning rates, one for each *a priori* cue value that cues could be assigned (Taswell et al., 2018). In addition, we utilized a novelty bonus parameter to better fit novelty-seeking behavior seen on the first trial where a novel option is introduced (Costa et al., 2014, 2019; Kakade & Dayan, 2002; Wittmann et al., 2008). We rescaled the prediction errors to be between −1 and 0 for negative prediction errors and 0 and 1 for positive prediction errors, to enable direct comparisons of learning rates across cues associated with different valences and outcome magnitudes. The parameters of all models used were optimized using the *fminsearch* function in MATLAB.

#### Novelty Bonus

This model had one term for choice consistency (β), four learning rates (α) – one for each *a priori* average cue value, and one novelty bonus term (γ). The novelty bonus term was a fixed constant, γ added to novel cues when they were introduced and estimated as a free parameter 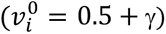. The model updates the value, *ν*, of a chosen option, *i*, based on choice feedback, *R,* in trial *t* as:

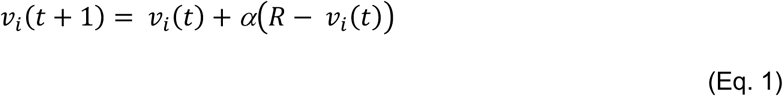

These value estimates were then passed through a softmax function to generate choice probabilities:

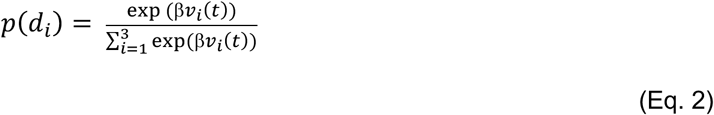

And then calculated the log-likelihood as:

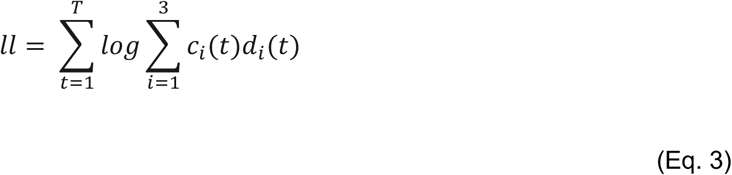

Where *c_i_*(*t*) = 1 when the subject chooses option *i* in trial *t* and *c_i_*(*t*) = 0 for all unchosen options. In other words, the model maximizes the choice probability, *d_i_*(*t*) of the actual choices the subjects made. *T* is the total number of trials in the session for each monkey. Parameters were maximized using standard techniques (Costa et al., 2014, 2019). To avoid local minima, initial value and learning rate parameters were drawn from a normal distribution with a mean of 0.5 and a standard deviation of 3. An alternative model fit two novelty bonuses, *γ^GAIN^* and *γ^LOSS^* that were fixed constants added to the value of the novel option when it was introduced based on whether selecting the best alternative option would result in a gain or a loss of tokens. Use of this model did not statistically improve model fit (*χ^2^* = 1.58, *p* = .2083).

#### Novelty Bonus and Perceptual Novelty Discount

To better fit the initial novelty preferences seen in the first few trials, which quickly falls off and asymptotes, we fit the same model described above but included a term that discounted perceptual novelty. This model had one term for choice consistency (β), four learning rates (α) – one for each *a priori* average cue value, one novelty bonus term (γ), and one term to discount the novel option once it was no longer perceptually novel to the monkey (*κ*). In this model, the first trial where a novel option was introduced assigned the cue value according to equation 1. The second trial where the novel option was selected, the value of the novel option was updated as follows:

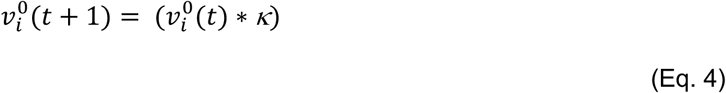

On all other trials, values were updated trial-by-trial according to equation 1, and then value estimates were passed through a softmax function to generate choice probabilities using equation 2. We performed a log likelihood ratio test and determined that the addition of the perceptual novelty discounting term provided a better fit than the model without it (*χ^2^* = 11.66, *p* = .0006). All data and code supporting the findings of this nonhuman primate study are available upon reasonable request from the corresponding author.

## Acknowledgements

This work was supported by funds provided by Oregon Health and Science University, Emory University, and National Institutes of Health: R01MH125824 (VDC), P51 OD011132 (VDC), and P51 OD011092 (VDC). We thank Drs. Craig Taswell and Diana Burke for helpful discussions about the use of conditioned reinforcers in the context of instrumental conditioning. We thank Drs. Vanessa Brown, Peter Hitchcock, and Jeremy Hogeveen for their helpful comments on the manuscript. Portions of this work were presented at the 2024 annual conference on Computational Cognitive Neuroscience.

## Author Contributions (using CRediT)

Conceptualization: VDC

Methodology: VDC, KMR, MDR, KH

Software: VDC, KMR

Formal analysis: KMR, VDC

Investigation: MDR, KH, VDC

Resources: VDC

Data curation: KMR, VDC, MDR

Writing-original draft: KMR, VDC

Writing-review and editing: KMR, VDC, MDR

Supervision: VDC

Project administration: VDC

Funding acquisition: VDC

